# Whole-brain comparison of rodent and human brains using spatial transcriptomics

**DOI:** 10.1101/2022.03.18.484766

**Authors:** Antoine Beauchamp, Yohan Yee, Ben Darwin, Armin Raznahan, Rogier B. Mars, Jason P. Lerch

## Abstract

The ever-increasing use of mouse models in preclinical neuroscience research calls for an improvement in the methods used to translate findings between mouse and human brains. Using openly accessible brain-wide transcriptomic data sets, we evaluated the similarity of mouse and human brain regions on the basis of homologous gene expression. Our results suggest that mouse-human homologous genes capture broad patterns of neuroanatomical organization, but that the resolution of cross-species correspondences can be improved using a novel supervised machine learning approach. Using this method, we demonstrate that sensorimotor subdivisions of the neocortex exhibit greater similarity between species, compared with supramodal sub-divisions, and that mouse isocortical regions separate into sensorimotor and supramodal clusters based on their similarity to human cortical regions. We also find that mouse and human striatal regions are strongly conserved, with the mouse caudoputamen exhibiting an equal degree of similarity to both the human caudate and putamen.

## Introduction

Animal models play an indispensable role in neuroscience research, not only for understanding disease and developing treatments, but also for obtaining data that cannot be obtained in the human. While numerous species have been used to model the human brain, the mouse has emerged as the most prominent of these, due to its rapid life cycle, straightforward husbandry, and amenability to genetic engineering *(1–4)*. Mouse models have proven to be extremely useful for understanding diverse features of the brain, from its molecular neurobiological properties to its large-scale network properties *(5–7)*. However, translating findings from the mouse to the human has not been straightforward. This is especially evident in the context of neuropsychopharmacology, where promising neuropsychiatric drugs have one of the highest failures rates in Phase III clinical trials *(8)*.

Successful translation requires an understanding of how effects on the brain of the model species are likely to manifest in the brain of the actual species of interest. This is not trivial in the case of the mouse and human, as the two species diverged from a common ancestor about 80 million years ago *(9)*. Although common themes are apparent in the brains of all mammalian species studied to date *(10)*, there remain substantial differences between the mouse and human brain. Beyond the obvious differences in size, large parts of the human cortex potentially have no corresponding homologues in the mouse *(11)*. Direct comparisons across the brains of different species are further complicated by the fact that researchers from different traditions use inconsistent nomenclature to refer to similar neuroanatomical areas *(12, 13)*.

Over the course of the last decade, we have developed novel approaches to explicitly evaluate similarities and differences between the brains of related species. These approaches describe brains using common data spaces that are directly comparable between species, making it possible to evaluate the similarity of different regions in a quantitative fashion *(14)*. This way, potential homologues can be formally tested and regions of the brain that do not allow for straightforward translation can be identified *(15)*. Establishing such a formal translation between the mouse and the human brain would allow scientists involved in translational research to explicitly test hypotheses about conservation of brain regions, identify regions that are well suited to translational paradigms, and directly transform quantitative maps from the brain of one species to the other.

One approach towards building these common spaces has been to exploit connectivity. The connections of a brain region tend to be unique and can therefore be seen as a diagnostic of an area *(16, 17)*. The common connectivity space approach relies on defining agreed upon neuroanatomical homologues a priori and then expressing the connectivity fingerprint of regions under investigation with those established homologues in the two brains *(18)*. The connections of any given region to the established homologues thus form a common space, which links the two brains. In a series of early studies, we compared the connectivity of the macaque and human brain, identifying homologies as well as specializations across association cortex *(19–21)*. The same approach has recently been applied to mouse-human comparisons for the first time, demonstrating conserved organization between the mouse and human striatum, but some specialization in the human caudate related to connectivity with the prefrontal cortex *(22)*. A similar study recently compared connectivity of the medial frontal cortex across rats, marmosets, and humans *(23)*. However, the lack of established neuroanatomical homologues in mice, particularly in the cortex, limits the use of connectivity to compare these species.

A more promising approach to mouse-human comparisons could be to exploit the spatial patterns of gene expression. Advances in transcriptomic mapping can be used to characterise the differential expression of many thousands of genes across the brain and compare the pattern between regions *(24)*. Moreover, the availability of whole-brain spatial transcriptomic data sets for multiple species provides an opportunity to run novel analyses at low cost *(25, 26)*. Such maps for the human cortex show topographic patterns that mimic those observed in other modalities, such as a gradient between primary and heteromodal areas of the neocortex *(27)*. Importantly, these patterns appear to be conserved across mammalian species *(28)*, which opens up the possibility of using the expression of homologous genes as a common space across species. In fact, a recent study demonstrated how the expression of homologous genes can be used to directly register mouse and vole brains into a common reference frame, which allows for direct point-by-point comparisons of brain maps *(29)*. However, this specific approach is only feasible because of the large degree of morphological similarity between mouse and vole brains. In the case of mouse-human comparisons, we almost certainly cannot directly register mouse and human brains into a common coordinate frame using methods for image registration. Hence we need to be more creative in our approach.

Here we examine the patterns of similarity between the mouse and human brain using a common space constructed from spatial gene expression data sets. We begin with an initial set of 2624 homologous genes. Subsequently, we present and evaluate a novel method for improving the resolution of mouse-human neuroanatomical correspondences using a supervised machine learning approach. Using the novel representation of the gene expression common space, i.e. a latent gene expression space, we analyse the similarity of mouse and human isocortical subdivisions and demonstrate that sensorimotor regions exhibit a higher degree of similarity than supramodal regions. Finally, we examine the patterns of transcriptomic similarity at a voxel-wise level in the mouse and human striatum.

## Results

### Homologous genes capture broad similarities in the mouse and human brains

We first examined the pattern of similarities that emerged when comparing mouse and human brain regions on the basis of their gene expression profiles. We constructed a gene expression common space using widely available data sets from the Allen Institute for Brain Science: the Allen Mouse Brain Atlas (AMBA) and the Allen Human Brain Atlas (AHBA) *(25, 26)*. These data sets provide whole-brain coverage of expression intensity for thousands of genes in the mouse and human genomes. For our purposes we filtered these gene sets to retain only mouse-human homologous genes using a list of orthologues obtained from the NCBI HomoloGene system (NCBI 2018). Prior to analysis, both data sets were pre-processed using a pipeline that included quality control checks, normalization procedures, and aggregation of the expression values under a set of atlas labels. The result was a gene-by-region matrix in either species, describing the normalized expression of 2624 homologous genes across 67 mouse regions and 88 human regions (see Materials and methods). We quantified the degree of similarity between all pairs of mouse and human regions using the Pearson correlation coefficient, resulting in a mouse-human similarity matrix (Fig. 1A).

**Fig. 1.**
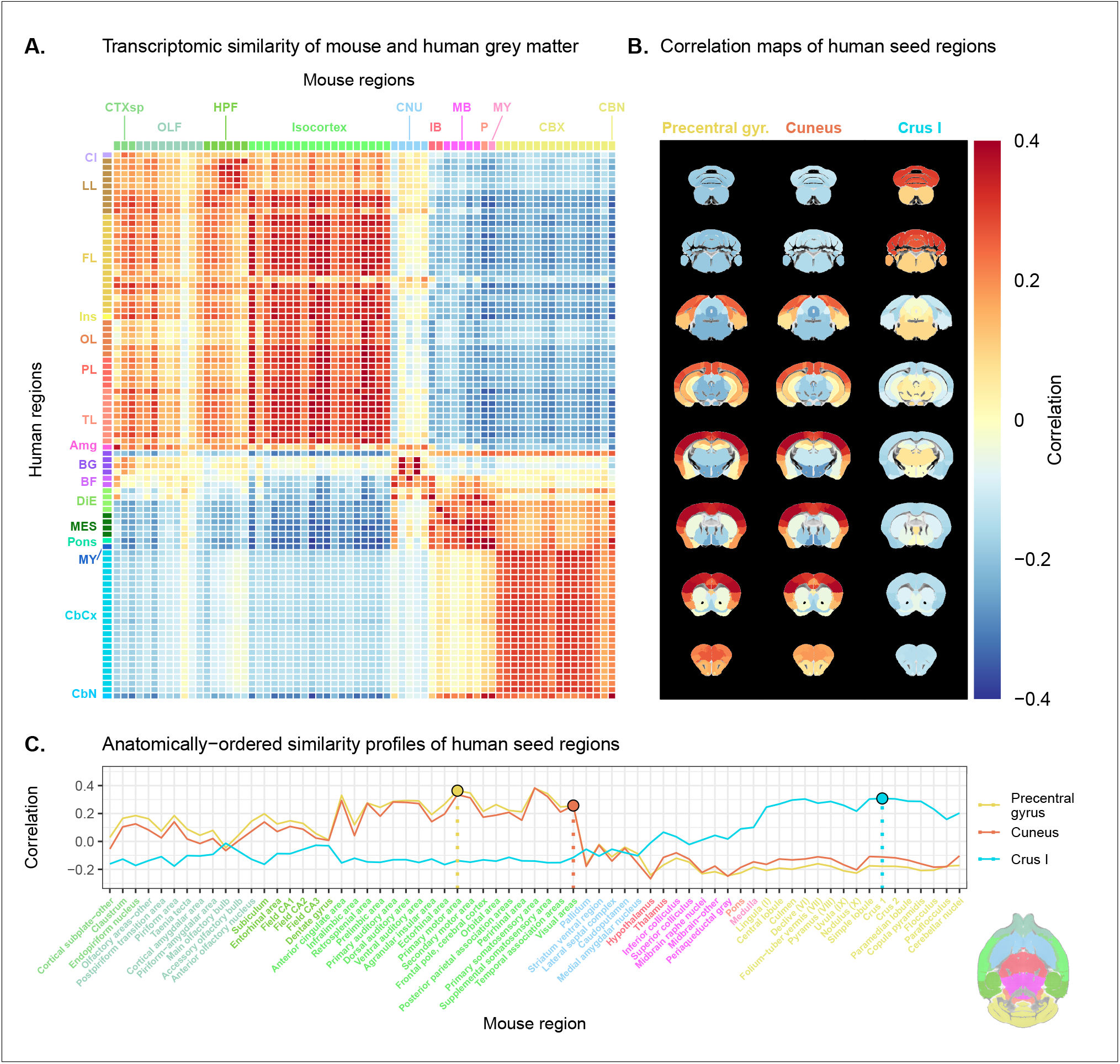
Transcriptomic similarity in the mouse and human brains. (**A**) Similarity matrix displaying the correlation between 67 mouse regions and 88 human regions based on the expression of 2624 homologous genes. Columns are annotated with 11 broad mouse regions: Cortical subplate (CTXsp), olfactory areas (OLF), hippocampal formation (HPF), isocortex, cerebral nuclei (CNU), interbrain (IB), midbrain (MB), pons (P), medulla (MY), cerebellar cortex (CBX), cerebellar nuclei (CBN). Rows are annotated with 16 broad human regions: Claustrum (Cl), limbic lobe (LL), frontal lobe (FL), insula (Ins), occipital lobe (OL), parietal lobe (PL), temporal lobe (TL), amygdala (Amg), basal ganglia (BG), basal forebrain (BF), diencephalon (DIE), mesencephalon (MES), pons, myelencephalon (MY), cerebellar cortex (CbCx), cerebellar nuclei (CbN). Broad patterns of similarity are evident between coarsely defined brain regions, while correlation patterns are mostly homogeneous within these regions. (**B**) Mouse brain coronal slices showing similarity profiles for the human precentral gyrus, cuneus and crus I. Correlation patterns for the precentral gyrus and cuneus are highly similar to one another and broadly similar to most isocortical regions. The crus I is homogeneously similar to the mouse cerebellum. (**C**) Anatomically-ordered line charts displaying the similarity profiles for the seed regions in (B). Dashed vertical lines indicate the canonical mouse homologue for each human seed. Annotation colours correspond to atlas colours from the AMBA and AHBA for mouse and human regions respectively.

We find that the similarity matrix exhibits broad patterns of positive correlation between the mouse and human brains. These clusters of similarity correspond to coarse neuroanatomical regions that are generally well-defined in both species. For instance, we observe that, overall, the mouse isocortex is similar to the human cerebral cortex, with the exception of the hippocampal formation, which forms a unique cluster. Similarly the mouse and human cerebellar hemispheres cluster together, while the cerebellar nuclei show relatively high correlation to each other (r = 0.404) as well as to brain stem structures like the pons (*r* = 0.359 and *r* = 0.371 for the mouse and human nuclei respectively) and myelencephalon (*r* = 0.318 and *r* = 0.374). The associations between broad regions such as these are self-evident in the correlation matrix.

Our ability to resolve regional matches on a finer scale is limited when using all homologous genes in this way. This is especially true for regions within the cerebral and cerebellar cortices, which exhibit a high degree of internal homogeneity. This is apparent in the similarity profiles, defined here as the set of correlation values between a given seed region and all target regions in the other species. For example, the human precentral gyrus and cuneus are most strongly correlated to many regions of the mouse isocortex. While the brain maps feature a rostral-caudal gradient (Fig. 1B), the profiles of the two seeds are highly similar despite the regions having very different functions (Fig. 1C). Indeed, the correlation between the similarity profiles of the precentral gyrus and cuneus is *r* = 0.980. The similarity profile of human cerebellar crus 1 highlights another example of this homogeneity. The profile of crus 1 is similar to that of all regions of the mouse cerebellum, with an average correlation of *r* = 0.269 and a standard deviation of *σ* = 0.041. Across all regions, the variance of the correlations across cortical regions is *σ*^2^ = 0.0052 while that across cerebellar hemispheric regions is *σ*^2^ = 0.0017, compared with a total variation of *σ*^2^ = 0.0416 across all entries in the matrix.

So, although these is some distinguishing power in the profiles of regions at a finer scale, this is much smaller than between coarse anatomical regions. This also true for parts of the broad anatomical systems that are part of the same functional system. This suggests that the regional expression patterns of mouse-human homologous genes can be used to identify general similarities between the brains of the two species even using a simple correlation measure, but the ability to identify finer scale matches might require a more subtle approach.

### A latent gene expression space improves the resolution of mouse-human associations

In the previous analyses, we showed that the expression profiles of homologous genes capture broad similarities across the mouse and the human for the major subdivisions of the brain. Some information at a finer resolution (e.g. within the isocortex) was also evident, but much less distinctive. Our next goal was to investigate whether it is possible to leverage the gene expression data sets to relate mouse and human brains to one another at a finer regional level. In order to do so, we sought to maximise the informational value in the set of 2624 homologous genes by creating a new latent common space that exploits the regional distinctiveness of the expression profiles.

The approach used in the previous analysis relied on using homologous genes as a common space between the mouse and human brain. This approach effectively assigns equal value to each gene, whereas a more powerful approach would be to weight genes by their ability to distinguish between different brain regions. We investigated whether we could accomplish this by constructing a new set of variables from combinations of the homologous genes. Our primary goal here was to transform the initial gene space into a new common space that would improve the locality of the matches. However while we sought a transformation that would allow us to recapitulate known mouse-human neuroanatomical homologues, we also wanted to avoid directly encoding such correspondences in the transformation. Using this information as part of the optimization process for the transformation would run the risk of driving the transformation towards mouse-human pairs that are already known. While we are interested in being able to recover such matches, we are equally interested in identifying novel and unexpected associations between neuroanatomical regions in the mouse and human brains (e.g. one-to-many correspondences). Given these criteria, our approach to identifying an appropriate transformation was to train a multi-layer perceptron classifier on the data from the AMBA. The classifier was tasked with predicting the 67 labels in our mouse atlas from the voxel-wise expression of the homologous genes (Fig. 2A).

**Fig. 2.**
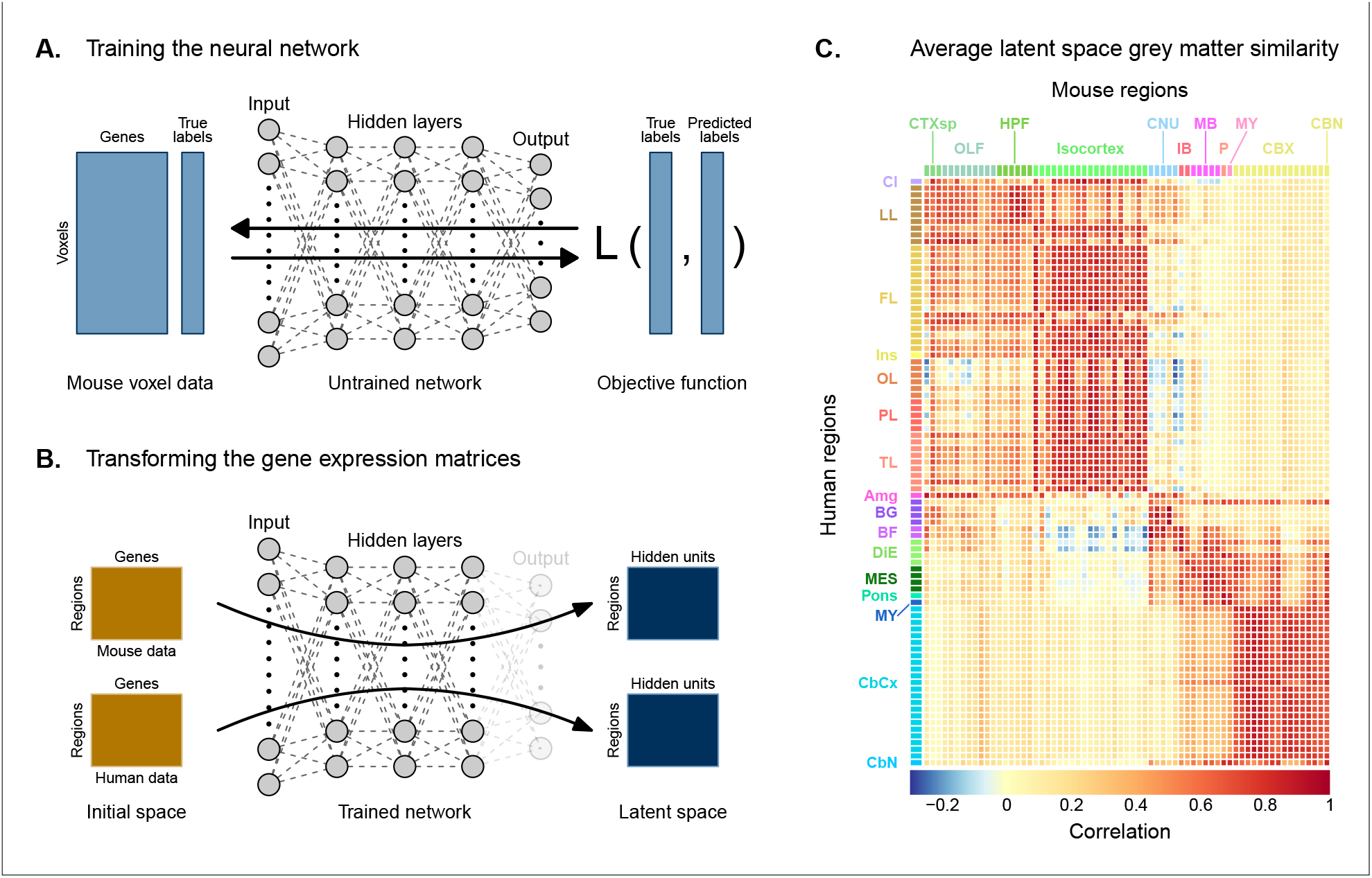
Creating a new common space. (**A**) Voxel-wise expression maps from 2624 homologous genes in the AMBA were used to train the neural network to classify each mouse voxel into one of 67 atlas regions. (**B**) Once the network is trained, the output layer is removed. The mouse and human regional gene expression matrices are passed through the network, resulting in lower-dimensional latent space representations of the data. The training and transformation process was repeated 500 times. (**C**) A similarity matrix displaying the gene expression latent space correlation between mouse and human regions, averaged over 500 neural network training runs. Similar brain regions exhibit very high correlation values. Column and row annotations as described in Figure 1.

While the model could have been trained using the data from either species, we chose to use the mouse data because it provides continuous coverage of the entire brain and is thus better suited to this purpose. In training the model to perform this classification task, we effectively optimize the network architecture to identify a transformation from the input gene space to a space that encodes information about the delineation between mouse brain regions. To extract this transformation, we removed the output layer from the trained neural network. The resulting architecture defines a transformation from the input space to a lower-dimensional gene expression latent space. We then applied this transformation to the mouse and human gene-by-region expression matrices to obtain representations of the data in the latent common space (Fig. 2B). Finally, we used these gene expression latent common space matrices to compute the new similarity matrix (Fig. 2C). Since the optimization algorithm used to train the perceptron features an inherent degree of stochasticity, we repeated this training and transformation process 500 times to generate a distribution of latent spaces and similarity matrices over training runs.

To assess whether the latent space representations of the data improved the resolution of the mouse-human matches, we considered two criteria. The first was whether the similarity profiles of the mouse atlas regions were more localized within the corresponding broad regions of interest (e.g. primary motor area within isocortex), compared with their similarity profiles in the original gene space. We term this the locality criterion. The second criterion was whether the degree of similarity between canonical neuroanatomical homologues improved in this new latent common space. We term this the homology criterion. The locality criterion tells us about our ability to extract finer-scale signal in these profiles, while the homology criterion informs us about our ability to recover expected matches in this finer-scale signal. To evaluate these criteria, we computed ranked similarity profiles for every region in the mouse brain, ordered such that a rank of 1 indicates the most similar human region. In addition, given the difference in absolute value between the input gene space and gene expression latent space correlations, we scaled the similarity profiles to the interval [0, 1] in order to make comparisons between the spaces.

We evaluated the locality criterion by examining the decay rate of the top of the similarity profiles. We reasoned that the plateau of similarity to a broad brain region, as seen in the anatomically-ordered similarity matrices and profiles (Fig. 1,A and C; Fig. 2C), would correspond to a similar plateau at the head of the rank-ordered profiles. Moreover, the emergence of local signal would manifest as an increase in the range between the peaks and troughs within the broad region. In the rank-ordered profiles, this would correspond to a faster rate of decay at the head of the profile. In order to quantify this decay, we computed the rank at which each region’s similarity profile decreased to a scaled value of 0.75. This was calculated for every mouse region in the initial gene space, as well as in each of the 500 gene expression latent spaces. As a measurement of performance between the two representations of the data, we then took the difference in this rank between each of the latent spaces and the original gene space (Fig. 3A). A negative rank difference indicates an improvement in the latent space.

**Fig. 3.**
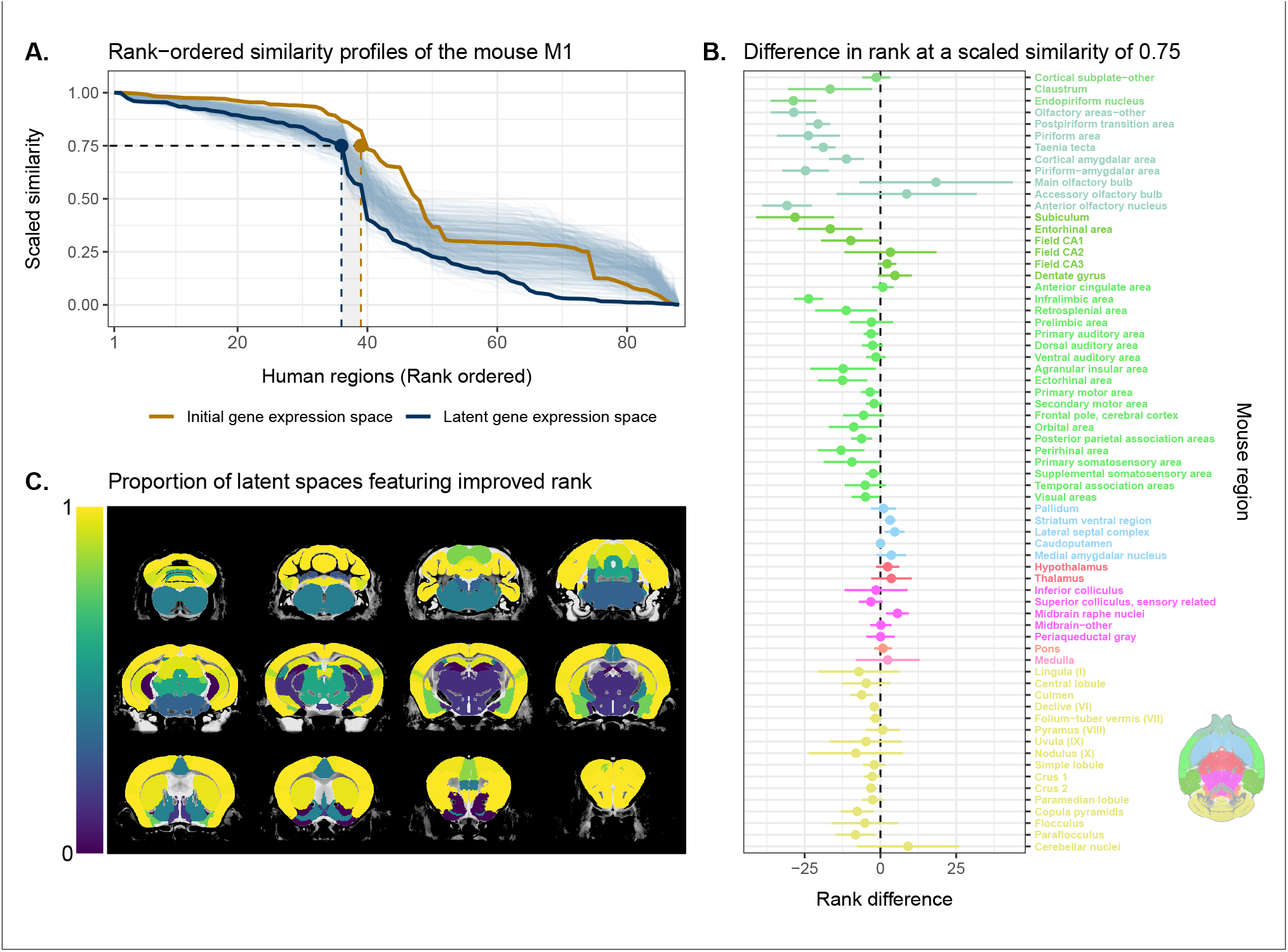
Quantifying improvement in locality in gene expression latent space. (**A**) The amount of local signal within a broadly similar region of the brain for a finer seed region’s (e.g. primary motor area) similarity profile can be quantified by the decay rate of the head of the rank-ordered profile. Decay rate was quantified by computing the rank at a similarity of 0.75. This metric was compared between the initial gene expression space (orange line) and every gene expression latent space resulting from repeated training of the neural network (every blue line is a training outcome, heavy blue line serves as an example). A negative difference between these rank metrics indicates an improvement in locality in the latent space. (**B**) Structure-wise distributions of differences in rank at a similarity of 0.75 between the initial gene expression space and the gene expression latent spaces. Points and error bars represent mean and 95% confidence interval. Dashed black line at 0 indicates the threshold for improvement in one space over the other. Colours correspond to AMBA annotations as in Figures 1 and 2. (**C**) Proportion of perceptron training runs resulting in an improvement or null difference in the gene expression latent space compared with the initial space. Cortical and cerebellar regions exhibit high proportions of improvement, while subcortical regions are less likely to be improved by the classification process.

Examining the structure-wise distributions of these rank differences, we found that for the majority of regions in our mouse atlas, the classification approach resulted in either an improvement in the amount of locality within a broad region, or no difference from the original gene space (Fig. 3, B and C). Specifically, 47 regions (70.1%) had a mean rank difference less than or equal to zero. Additionally, the same number of regions returned non-positive rank differences in at least 80% of latent spaces. A few regions performed considerably worse in the latent spaces, notably the main olfactory bulb (*µ* = 18.4; *σ* = 12.7), the accessory olfactory bulb (*µ* = 8.7; *σ* = 11.6), and the cerebellar nuclei(*µ* = 9.1; *σ* = 8.5). In particular, the main olfactory bulb performed worse in 96.6% of latent spaces. Regions within the cortical subplate and olfactory areas (e.g. endopiriform nucleus, postpiriform transition area) benefited the most from the classification approach, with many regions showing improvements in all latent spaces. While the effects are smaller, the similarity profiles of regions belonging to the isocortex and cerebellar cortex also saw an improvement in locality. In the isocortex, 16 out of 19 regions (84.2%) improved in at least 96% of latent spaces. In the cerebellar cortex, 73.3% of regions saw a similar improvement. In contrast, regions belonging to the cerebral nuclei, the diencephalon, midbrain and hindbrain did not see much improvement in this new common space. For instance, only 13.2% of latent spaces returned a non-positive rank difference in the thalamus. For many such regions the degree of locality appears to be worse in this space, though only by a small number of ranks (e.g. striatum ventral region, thalamus, midbrain raphe nuclei). Indeed, the mean rank difference and standard deviation over these regions and all latent spaces are *µ* = 1.4 and *σ* = 3.6. These results demonstrate that the supervised learning approach used here can improve the resolution of neuroanatomical correspondences between the mouse and human brains, though the amount of improvement varies over the brain. Regions that were already well-characterized using the initial set of homologous genes (e.g. subcortical regions) did not benefit tremendously, but numerous regions in the cortical plate and subplate, as well as the cerebellum, saw an improvement in locality in this new common space.

While the supervised learning approach improved our ability to identify matches on a finer scale for a number of brain regions, this does not necessarily mean that those improved matches are biologically meaningful. The second criterion for evaluating the performance of the neural network addresses whether this improvement in locality captures what we would expect in terms of known mouse-human homologies. To this end, we examined the degree of similarity between established mouse-human neuroanatomical pairs, both in the initial gene expression space and in the set of latent spaces. We began by establishing a list of 36 canonical mouse-human homologous pairs on the basis of common neuroanatomical labels in our atlases. For each of these regions in the mouse brain, we compared the rank of the canonical human match in the rank-ordered similarity profiles between the latent spaces and the original gene expression space (Fig. 4A). The lower the rank, the more similar the canonical pair, with a rank of 1 indicating maximal similarity. We additionally calculated the proportion of latent spaces in which each mouse region was more similar or as similar to its canonical human match compared with the initial gene space (Fig. 4B).

**Fig. 4.**
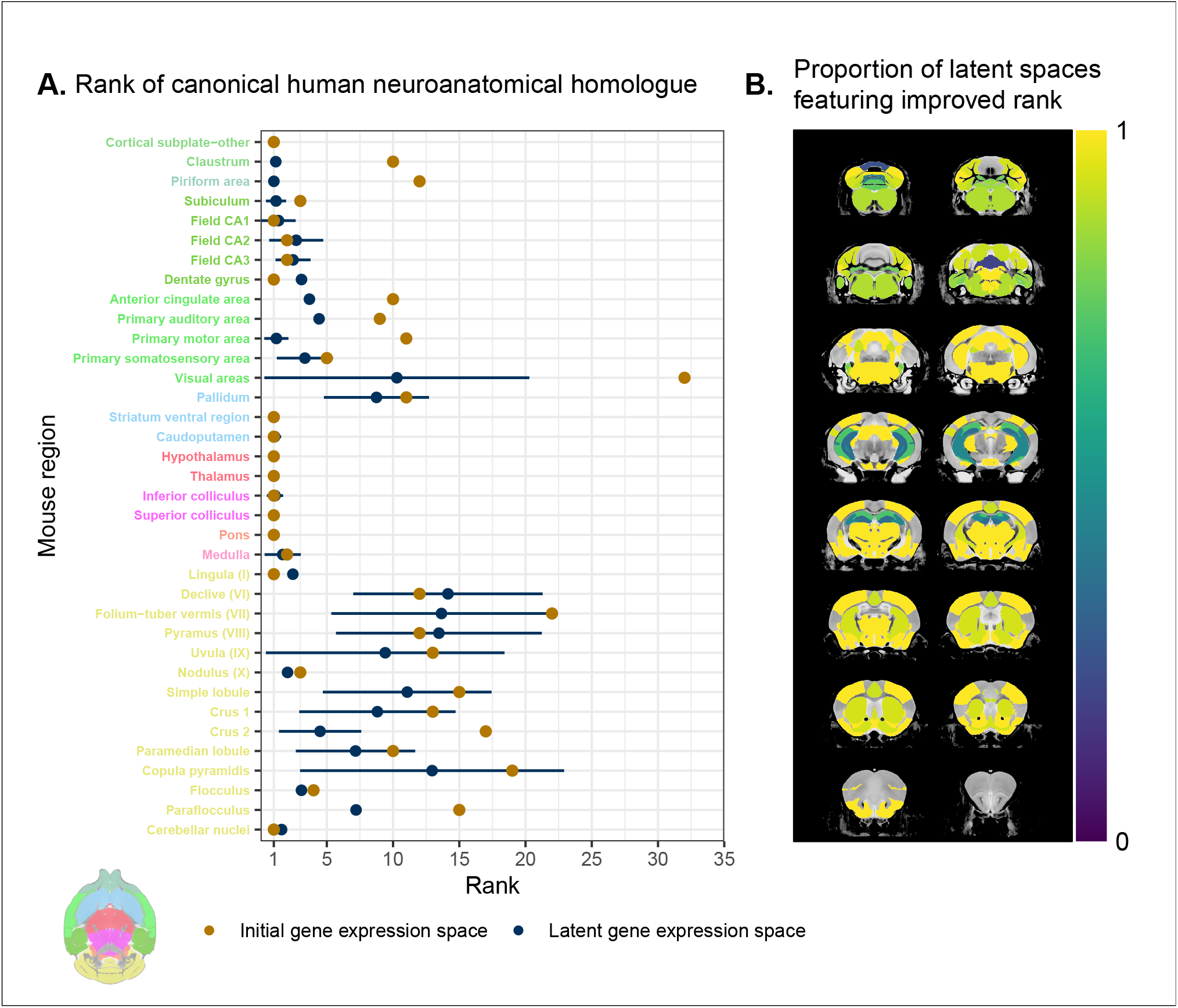
Recovering canonical neuroanatomical pairs in gene expression space. (**A**) Comparison between the ranks of canonical human matches for mouse seed regions between the initial gene expression space and gene expression latent spaces. Points and error bars represent mean and 95% confidence interval. Mouse region names are coloured according to the AMBA palette. (**B**) Proportion of latent spaces resulting in an improvement or null difference compared with the initial gene space. Uncoloured voxels correspond to regions with no established canonical human match.

We find that for most regions in this subset, the classification approach either improves the correspondence or performs as well as the full set of homologous genes. For example, 73% of regions exhibit improved similarity in at least 80% of latent spaces. The improvement is most pronounced for regions in the cortical subplate and isocortex. In particular, the visual areas improve from a rank of 32 to an average of 10, though the variance is much higher in this case. Many regions in the sub-cortex do not benefit from the gene expression latent spaces since the initial gene set was already recapitulating the appropriate match with maximal similarity. Apart from the pallidum and the medulla, each of these regions is maximally similar to its canonical match in at least 90% of latent spaces. In such cases, the classification approach performs as well as the original approach. Finally, although many regions in the cerebellum feature some improvement in the latent spaces, the variation in the rank of the standard human pair is often quite large, indicating some instability in the neural network’s ability to recover these matches. However, while the rank of the canonical pair varies in different instances of the latent space, the top matches for any given cerebellar region are always cerebellar regions. For instance, when the mouse crus 1 is used as the seed region, the human crus 1 is most often assigned a rank between 6 and 9. However, similar proportions in that range occur for the crus 2 and lobules V, VI and VIIB, indicating that these cerebellar regions are swapping ranks in the different latent spaces. Thus while cerebellar regions are reliably associated with other cerebellar regions in the gene expression latent spaces, these associations are not stable over multiple training runs.

Together, these results demonstrate that the multi-layer peceptron classification approach improves our ability to resolve finer scale mouse-human neuroanatomical matches within the broadly similar regions obtained using the initial gene expression space. By training a classifier to predict the atlas labels in one species, we were able to generate a new common space that amplified the amount of local signal within broadly similar regions while also improving our ability to recover known mouse-human neuroanatomical pairs.

### Cortical areas involved in sensorimotor processing show greater transcriptomic similarity than supramodal areas

It is well established that the brains of most, if not all, extant mammalian species follow a common organizational blueprint inherited from an early mammalian ancestor *(30)*. A number of cortical subdivisions have consistently been identified in members of many distantly related mammalian species *(10)* and hypothesized to have been present in the common ancestor of all mammals *(30)*. While it is clear that basic sensorimotor cortical regions are found in the majority of mammals, including mice and humans, there is much debate about the extent to which cortical areas involved in supramodal processing are conserved across mammalian taxa. Although some supramodal regions were likely present in the earliest mammals, including some cingulate regions and an orbitofrontal cortex *(30)*, since the divergence of mouse and human lineages some 80 million years ago, the primate neocortex has undergone substantial expansion and re-organization *(9)*. Indeed, when comparing the human neocortex even to primate model species, this is the likely locus of areas that cannot be easily translated between species *(15)*. As a result, it is important to investigate whether our between-species mapping is more successful in somatosensory areas than supramodal areas.

We assessed the similarity between mouse and human isocortical areas using the pairwise correlations in each of the gene expression latent spaces returned from the multi-layer perceptron. For every region in the mouse isocortex, we evaluated the distribution of maximal correlation values over latent spaces (Fig. 5A). While the region-wise variance for each isocortical area was large, we found that, on average, sensorimotor regions exhibited higher maximal correlation values than supramodal regions. The mouse primary somatosensory and motor areas have the highest average maximal correlation values, with *r* = 0.94*±*0.04 and *r* = 0.93*±*0.04 respectively. We additionally examined the distributions of maximal correlation, grouped by cortex type (Fig. 5B). To generate these distributions, we computed average maximal correlation values by cortex type in each of the latent spaces. Here too we find that that sensorimotor regions are associated with higher maximal correlation values on average (*r* = 0.89 *±* 0.04), compared with supramodal areas (*r* = 0.85 *±* 0.03). These distributions demonstrate that sensorimotor isocortical regions exhibit more similarity overall on the basis of homologous gene expression than do supramodal regions.

**Fig. 5.**
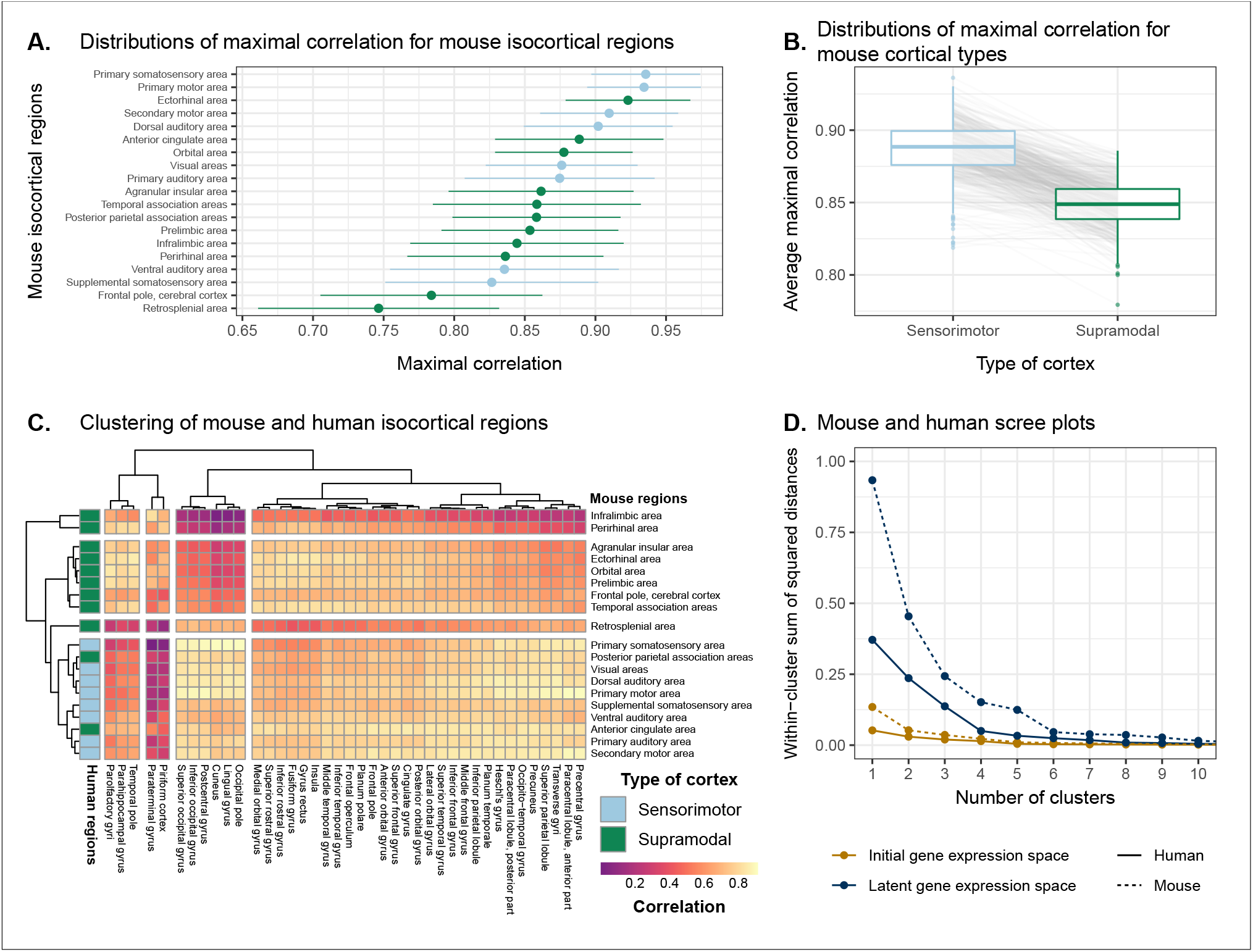
Similarity of mouse-human isocortical regions. (**A**) Maximal correlation distributions of mouse isocortical regions. Points and error bars represent mean and 95% confidence interval over latent space samples. (**B**) Distributions of average maximal correlation for sensorimotor and supramodal isocortical areas in each gene expression latent space. Grey lines correspond to individual latent spaces. (**C**) Hierarchical clustering of mouse and human isocortical regions based on average latent space correlation values. Mouse regions are annotated as sensorimotor or supramodal. Four clusters were chosen for visualization using the elbow method. (**D**) Within-cluster sum of squared distances for different numbers of mouse and human isocortical clusters in the average latent space and initial homologous gene space.

While we found that sensorimotor isocortical areas in the mouse brain were more similar to human brain regions than supramodal areas, the distributions of maximal correlation do not speak to the neuroanatomical patterns of organization for these matches. To understand how the similarity patterns of mouse and human cortical subdivisions were organized, we used hierarchical clustering to cluster mouse and human isocortical regions on the basis of their similarity profiles in the average gene expression latent space (Fig. 5C). This allows us to examine the similarity of regions to one another within and across brains at multiple levels simultaneously.

At a high level, we find a striking segregation of the mouse isocortex into one main cluster that corresponds to regions that are primarily engaged in sensorimotor processing and separate clusters of regions that are supramodal. All of the sensorimotor areas cluster together, but three supramodal areas also form part of this cluster: the retrosplenial area, the posterior parietal association areas, and the anterior cingulate cortex. Of these, the retrosplenial area is the most different, being the first to separate out from the other regions. In fact, the retrosplenial area is the mouse isocortical region with the smallest correlation values (Fig. 5A). The mouse sensorimotor cluster is characterized by high correlation values to human sensorimotor regions like the precentral gyrus, the cuneus, and the postcentral gyrus, as well as low correlation values to the piriform cortex and paraterminal gyrus.

At this level of clustering, the remaining mouse supramodal subdivisions form two clusters. These both exhibit low similarity to the human somatosensory and visual areas, but the cluster containing the infralimbic and perirhinal areas additionally exhibits low correlation values with the precentral gyrus, anterior paracentral lobule, and transverse gyri. The human cortical regions do not segregate as cleanly into sensorimotor and supramodal clusters. Under a similar level of description of four clusters of areas, the majority of areas belong to a large cluster that includes a mix of cortical types. However, at a lower level of the hierarchy, if the number of clusters is increased to five, this large cluster breaks up into two smaller clusters that feature some delineation between supramodal and sensorimotor areas, which are primarily motor and auditory in nature (e.g. precentral gyrus, Heschl’s gyrus). Interestingly, the postcentral gyrus, i.e. primary somatosensory area, forms a separate cluster with a set of visual areas such as the cuneus and lingual gyrus. These regions exhibit very similar correlation profiles to the mouse isocortical regions, including maximal correlation to the mouse primary somatosensory area, with an average of *r* = 0.92. The cluster is characterized by high correlations to the mouse sensorimotor cluster and low correlations to the mouse supramodal clusters. Overall the human sensorimotor isocortical regions are loosely organized in clusters that contain sensory-visual areas and auditory-motor areas.

We additionally ran hierarchical clustering on the isocortical similarity matrix in the original homologous gene space. While the cluster annotations were not substantially different in this space, we observed that the Euclidean distances within and between clusters were smaller compared with the latent space clustering, further confirming that the perceptron classification approach improves the segregation of brain regions in the gene expression common space (Fig. 5D).

Overall, we observe a greater degree of similarity between mouse and human cortical regions involved in basic sensorimotor processing compared with supramodal or association areas. This is in line with the large body of existing research that suggests that sensory and motor areas of the cortex are conserved across the brains of mammals. While sensorimotor areas exhibit a greater degree of similarity than supramodal areas, the neuroanatomical pattern of correspondences obtained using mouse-human homologous genes is not at the level of individual cortical areas. Still, using a clustering approach we identified clear distinctions in the patterns of similarity between sensorimotor and supramodal areas, especially for regions in the mouse isocortex.

### Transcriptomic comparison of the mouse and human striatum

We have focused here on comparing mouse and human brain organization using transcriptomic data, with a latent space based on homologous genes as the common space between the two species. To date, common space comparisons between the mouse and human brain have only been performed using functional connectivity *(22, 23)*. As a case in point, Balsters et al. (2020) compared mouse and human striatal organization using this measure. They found that the nucleus accumbens was highly conserved between mice and humans, and that voxels in the posterior part of the human putamen were significantly similar to the lateral portion of their mouse caudoputamen parcellation. Additionally, they report that 85% of voxels in the human striatum were not significantly similar to any of their mouse striatal seeds, and that 25% of human striatal voxels were significantly dissimilar compared with the mouse. These differences were understandable, as they involved parts of the human striatum that connected to parts of prefrontal cortex that have no known homolog in the mouse *(21)*. However, it is not necessarily the case that between-species differences in connectivity are associated with distinct architectonic or molecular signatures. Therefore, we investigated the patterns of similarity between the mouse and human striata on the basis of gene expression using the neural network latent space representations.

We first identified the striatal regions present in the Allen human dataset: the caudate, the putamen, and the nucleus accumbens. We evaluated the correlation between the microarray samples in these regions and every region in the mouse atlas. Based on these correlation values, we focused our analysis on the four mouse regions that were consistently the most similar across all latent spaces: the caudoputamen, the nucleus accumbens, the fundus of striatum, and the olfactory tubercle. For each of the human striatal regions, we then calculated the average correlation over the samples to each of the mouse targets. We examined the distribution of these average correlation values over the latent spaces (Fig. 6A). We find that the human caudate and putamen consistently exhibit the strongest degree of similarity to the mouse caudoputamen. The median of the distribution for the caudate-caudoputamen pairs is 0.95 and 0.97 for the putamen-caudoputamen pairs, with modal values of 0.94 and 0.97, respectively. All latent spaces return correlations greater than 0.9 for caudate-caudoputamen and putamen-caudoputamen pairs. Beyond this expected top match, the caudate and putamen both exhibit high similarity to the nucleus accumbens and the fundus of striatum, with mean correlation values of about 0.80. Neither of these target regions is consistently more similar to the mouse caudoputamen over all latent spaces.

**Fig. 6.**
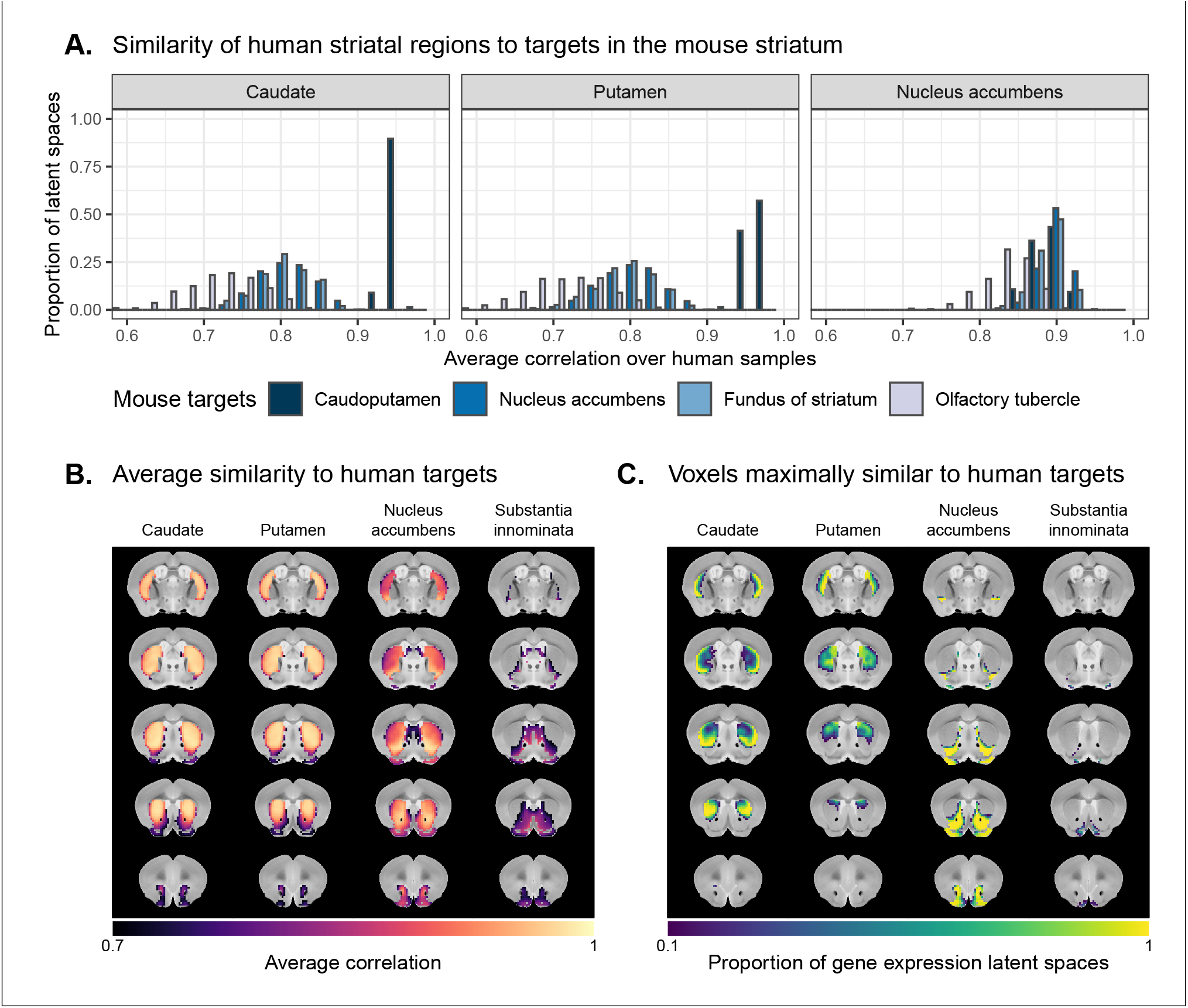
Similarity among mouse and human striatal regions. (**A**) Distributions over gene expression latent spaces of region-wise average correlation values for mouse and human striatal pairs. Human regions were chosen based on the AHBA ontology. Mouse target regions were chosen to be those with the highest average correlation values. (**B**) Latent space averaged correlations between voxels in the mouse striatum and human target regions. Target regions were selected based on the highest mean correlation across all striatal voxels. (**C**) Proportions of latent spaces in which mouse striatal voxels are maximally similar to human target regions.

While the similarity of the caudate and the putamen to the caudoputamen is unsurprising, the story is not as clear for the human nucleus accumbens. We find that the variance in correlation calculated over all mouse targets is much lower (*σ* = 0.04) compared with the equivalent variances for the caudate (*σ* = 0.09) and putamen (*σ* = 0.10), indicating less specificity to any one mouse striatal target. In particular, the human nucleus accumbens isn’t as specifically similar to the mouse nucleus accumbens in the way that the caudate and putamen are similar to the caudoputamen. The mouse target distributions are right-shifted compared with those for the caudate and putamen, with mean values of 0.90, 0.89, and 0.89 for the mouse nucleus accumbens, caudoputamen, and fundus of striatum, respectively. The human accumbens also exhibits a high degree of similarity to the mouse olfactory tubercle, the distribution of which is also right-shifted compared with the caudate and putamen.

Given the high correlation of the human caudate and putamen to the mouse caudoputamen, as well as the finding reported by Balsters et al. about the similarity of the lateral caudoputamen to the putamen, we were curious as to whether we could identify sub-regional patterns of similarity in the caudoputamen and other striatal regions using these gene expression data. To probe this question, we first examined the average latent space correlation between each voxel in the mouse striatum and every region in the human atlas. We created brain maps for the human regions that exhibited the highest mean correlation values, averaged over mouse striatal voxels: the caudate, the putamen, the nucleus accumbens, and the septal nuclei (Fig. 6B). We find that voxels in the caudoputamen exhibit a homogeneous pattern of similarity to both the caudate and the putamen. On average, voxels in the caudoputamen have a correlation of 0.95 to the caudate and 0.94 to the putamen, with standard deviations of 0.04 and 0.05 respectively. The caudate and putamen are associated with correlations of at least 0.90 in 88% and 84% of caudoputamen voxels. A number of voxels are also highly similar to the human nucleus accumbens, with an average correlation value of 0.90 and 55% of voxels returning a correlation of at least 0.9. The caudoputamen voxels most similar to the nucleus accumbens lie in the ventral-rostral part of the region. Of course, voxels in the mouse nucleus accumbens are also highly similar to the human nucleus accumbens, with an average of 0.90 and standard deviation of 0.06. While the human nucleus accumbens is the most strongly correlated region, a number of voxels also exhibit reasonably strong correlations to the substantia innominata, the septal nuclei, and the amygdala. Indeed, 91% of voxels in the accumbens are correlated at a value of 0.7 or higher to the substantia innominata. The equivalent percentages for the septal nuclei and amygdala are 78% and 74% respectively.

We additionally examined the proportion of latent spaces in which each voxel in the mouse striatum was maximally similar to the human target regions (Fig. 6C). As expected, we find that voxels in the caudoputamen are most often maximally similar to the human caudate and putamen, with 75% of voxels in the caudoputamen being maximally similar to the caudate or putamen in at least 95% of latent spaces, and 62% of voxels being maximally similar to one of those targets in all latent spaces. Interestingly, we observe the emergence of a continuous bilateral pattern of specifity to the caudate and putamen, with voxels in the rostral and lateral-caudal parts of the caudoputamen being more specific to the caudate, and voxels in the medial-rostral part being more specific to the putamen. This map highlights subtle differences in the similarity between caudoputamen voxels and the caudate or putamen. While this pattern distinguishes the two regions on the basis of which is the top match, individual voxels have very similar correlation values to the targets (Fig. 6B), with a mean difference in correlation of only 0.006. Beyond the caudoputamen, we find that the accumbens and olfactory tubercle in the mouse are consistently similar to the human nucleus accumbens, with 80% of mouse accumbens voxels and 65% of olfactory tubercle voxels having the human accumbens as their top match in at least 80% of latent spaces. For those voxels below this threshold, the human regions that are most often the top match are once again the amygdala, the septal nuclei, and the substantia innominata.

Overall, we observe a strong association between the mouse caudoputamen and both the human caudate and putamen. While we find a subtle pattern of specificity to either region among voxels in the caudoputamen on the basis of maximal similarity, the high degree of similarity in the correlation values to each region suggests that the majority of voxels in the caudoputamen are equally similar to the caudate and the putamen on the basis of the expression of mouse-human homologous genes. We also find that the nucleus accumbens is well conserved across species. However the region also exhibits patterns of similarity that go beyond the simple one-to-one match. The human accumbens features similar correlation values to the mouse caudoputamen and fundus of striatum, in addition to the accumbens proper, with no sharp distinction between these regions. It also exhibits a larger degree of similarity to the mouse olfactory tubercle. This is also seen in the mouse striatum, where voxels in the accumbens and the olfactory tubercle map onto the human accumbens.

## Discussion

We have demonstrated how spatial transcriptomic patterns of homologous genes can be used to make quantitative comparisons between the mouse and human brain. We showed that using homologous genes as a common space allows one to easily identify coarse similarities in brain structures across species, but that more fine-scaled parcellations, such as at the level of cortical areas, are more complex. Despite this limitation, the approach still allows for a formal assessment of different patterns of between-species similarity in primary compared to supramodal regions, identifications of distinct clusters of cortical territories across species, and comparison of between-species similarities at the transcriptomic level to those observed using other modalities. We will discuss our observations in the context of the importance of the mouse as a model for human neuroscience below.

The abundance of neuroscience research performed using mice has resulted in a wealth of knowledge about the mouse brain. In the preclinical setting, mouse models are utilized with the intention of better understanding human neuropathology. For instance, in the context of autism spectrum disorders, a plethora of studies using mouse models have reported on the neurobiological and neuroanatomical phenotypes that arise from mutations at specific genetic loci *(31–33)*. It is common for researchers involved in translational neuroscience to rely on findings of this kind to make inferences about the human disorder. The typical approach, which is to identify rough post-hoc correspondences between neuroanatomical ontologies, is not particularly comprehensive and is subject to confirmation bias. While it may be a reasonable starting point for comparison, the true correspondence between the mouse and human brain is likely more complicated given the evolutionary distance between the two species. Although overall patterns of brain organization, including the general pattern of neocortical organization, are similar across all mammals, substantial differences are evident *(34)*. To make matters worse, researchers from the different neuroscientific traditions often use distinct terminology, further complicating detailed information exchange. To address these problems, we sought to establish a first quantitative whole-brain comparison between the two species.

The expression of homologous genes provides an elegant way to define a common space for quantitative cross-species comparisons since it relies on homology at a deep molecular biological level. The approach is not without limitations however. First, the acquisition of whole-brain transcriptomic data is labour-intensive, time consuming, and invasive. These data sets cannot be generated easily, especially in the human, in which the process depends on the availability of post-mortem samples. As a result, the effective sample sizes are extremely limited in this domain. For instance, in the Allen mouse coronal in-situ hybridization data set used here, the brain-wide expression of each gene is sampled only once (barring a few exceptions). This constrains the types of analyses that are possible (e.g. null hypothesis significance testing) and largely limits the availability of replication data sets. That being said, new technologies, such as spatial transcriptomics, are gradually making it easier to acquire brain-wide gene expression data in less time and at lower cost *(24, 35, 36)*. Second, the approach of relying on all available genes is subject to noise. To address this issue, Myers (2017) *(37)* used a method of gene set selection to attempt to improve the correspondence between established mouse-human homologies. While this lead to improvement, it was only at the level of coarsely defined regions (e.g. cortex-cortex). Our approach, therefore, was to use supervised machine learning to create a latent common space based on combinations of homologous genes that can delineate areas within a single species.

This latent common space approach led to a substantial improvement in specificity of between-species comparisons. Nevertheless, it is evident that the first major distinction in gene expression patterns within a species, and the easiest identification of similarity across species, is at the coarse anatomical level of the major subdivisions of the vertebrate brain, such as the isocortex, cerebellar hemispheres and nuclei, and brain stem. All of these territories were present in the ancestral vertebrate brain *(38)* and the ability to detect conserved transcriptomic signatures at this level is not surprising. Within such structures, such as the isocortex, our ability to make simple one-to-one correspondences decreased. This is partly because areas within a coarse structure have more similar transcriptomic profiles, but also likely due to the fact that a single area in one brain does not have a single correspondent in another, larger brain. In other words, regions in the brains of related species may exhibit one-to-many or many-to-many mappings. In our study, we found greater cross-species similarity between cortical areas associated with sensorimotor processing than areas in supramodal cortex. Primary areas, including the sensorimotor areas, are present in all mammals studied to date and likely part of the common ancestors of all mammals *(10, 30)*. Although this common ancestor likely also had non-primary areas, it cannot be denied that association cortex expanded dramatically in primates and especially so in the human brain *(39, 40)*. Again, the pattern found here of greater similarity in more conserved areas might reflect this evolutionary history. In that context it is interesting to note that some non-primary areas thought to be present in the common mammalian ancestor, such as cingulate and orbitofrontal cortex *(30)* showed relatively high correlation to human areas.

An advantage of the approach presented here is that it can, in principle, be applied to any aspect of brain organization. Beyond simply establishing whether areas are similar across species in a particular common space, comparing the results across common spaces established using different types of neuronal data can inform on which larger principles of organization are similar across brains *(41)*. This is illustrated here by the results of our striatal analysis. We found high similarity between the human caudate and putamen and mouse caudoputamen, with little differentiation within these regions in a single species. In contrast, Balsters et al. (2020) demonstrated that human caudoputamen contains a distinct pattern of connectivity. At first sight, one could argue the results are in contrast. However, evolutionarily speaking, it is quite probable that an overall similar transcriptomic signature of the striatum can be accompanied by a distinct connectivity pattern to areas of the cortex present in only one of the two species. Indeed, this speaks to the different types of similarity that can be studied, depending on which aspect of brain organization one is interested in. Although the human brain is much larger than the mouse brain and contains a number of cortical territories that have no homologue in the mouse brain *(42, 43)*, the similarity in transcriptomic signature mean that translations between the species is valid in many contexts.

The power of a formal understanding of similarities and differences between brains at different levels of organization is evident. In fundamental neuroscience, it will help translate results from data types that cannot be obtained in humans to the human brain *(44)*. In translational neuroscience, it will, in a negative sense, help establish the limits of the translational paradigm by showing which aspects of the human brain cannot be understood using the model species *(45)*. In a positive sense, it will also help by establishing and improving our understanding of the many aspects in which the model and human brain do concur *(46)*. More ambitious still, it can provide a way in which highly diverse manifestations of certain disease syndromes (e.g. autism spectrum disorder) *(47, 48)* and the availability of many distinct model strains *(49)*, each hypothesized to capture a distinct aspect of a multi-dimensional clinical syndrome, can be related to one another. Ultimately, we believe that using the mapping of homologous gene expression between species can be an important part of building a transform that maps information obtained using mice to humans and vice versa.

## Materials and methods

### Mouse gene expression data

We used the adult mouse whole-brain in-situ hybridization data sets from the Allen Mouse Brain Atlas *(25)*. Specifically, we used 3D expression grid data, i.e. expression data aligned to the Allen Mouse Brain Common Coordinate Framework (CCFv3)*(50)* and summarized under a grid at a resolution of 200*µm*. We downloaded the gene expression “energy” volumes from both the coronal and sagittal in-situ hybridization experiments as a sequence of 32-bit float values using the Allen Institute’s API (http://help.brain-map.org/display/api/Downloading+3-D+Expression+Grid+Data). These volumes were subsequently reshaped into 3D images in the MINC format. Origin, extents, and spacing were defined such that the image was RAS-oriented, with the origin at the point where the anterior commissure crosses the midline. The MINC images from the coronal and sagittal data sets were then processed separately using the Python programming language. The sagittal data set was first filtered to keep only those genes that were also present in the coronal set. Images were imported using the pyminc package, masked and reshaped to form an experiment-by-voxel expression matrix. We pre-processed this data by first applying a log2 transformation for consistency with the human data set. For those genes associated with more than one in-situ hybridization experiment, we averaged the expression of each voxel across the experiments. We subsequently filtered out genes for which more than 20% of voxels contained missing values. Finally, we applied a K-nearest neighbours algorithm to impute the remaining missing values. The result of this pre-processing pipeline was a gene-by-voxel expression matrix with 3958 genes and 61315 voxels for the coronal data set and a matrix with 3619 genes and 26317 voxels for the sagittal data set.

### Human gene expression data

Human gene expression data was obtained from the Allen Human Brain Atlas *(26)*. The data were downloaded from the Allen Institute’s API (http://api.brain-map.org) using the abagen package in Python (https://abagen.readthedocs.io/en/stable/) *(51)*. We used the microarray data from the brains of all six donors, each of which contains log2 expression values for 58692 gene probes across numerous tissue samples. The data were pre-processed using a custom pipeline built following the recommendations from Arnatke-viciūtė et al. *(52)*. The pipeline was implemented using the R programming language. Specifically, once imported, we passed the data from individual donors through a set of filters. The first filter removed gene probes that were not associated with an existing Entrez gene ID. The second filtering step used the probe intensity filter provided by the AHBA. For each donor, we only retained the probes for which more than 50% of samples passed the intensity filter. After filtering, we aggregated the expression values for probes that corresponded to the same gene. To do so, we computed the average expression per sample for probes corresponding to a given gene. This was done separately for each donor, and the averages were computed in linear space rather than log2 space. Once the average gene expression values were obtained, we transformed the data back to log2 space. Finally, we combined the gene-by-sample expression matrices across the different donors. In doing so, we retained only those genes present in the data sets from all six donors. The result was a gene-by-sample expression matrix with 15125 genes and 3702 samples.

### Mouse atlases

We used a version of the DSURQE atlas from the Mouse Imaging Centre *(53–57)*, modified using the AMBA hierarchical ontology, which was downloaded from the Allen Institute’s API. The labels of the DSURQE atlas correspond to the leaf node regions in the AMBA ontology, which allowed us to use the hierarchical neuroanatomical tree to aggregate and prune the atlas labels to the desired level of granularity. For the purposes of our analyses, we removed white matter and ventricular regions entirely. The remaining grey matter regions were aggregated up the hierarchy so that the majority of resulting labels contained enough voxels to be classified appropriately by the multi-layer perceptron. In doing so, we maintained approximately the same level of tree depth within a broad region (e.g. cerebellar regions were chosen at the same level of granularity). This resulted in a mouse atlas with 67 grey matter regions. We additionally generated an atlas with 11 broader regions for visualization and annotation purposes.

### Human atlases

We used the hierarchical ontology from the AHBA, which we obtained using the Allen Institute’s API. We aggregated and pruned the neuroanatomical hierarchy to correspond roughly to the level of granularity obtained in our mouse atlas, resulting in 88 human brain regions. We additionally generated a set of 16 broad regions for visualization and annotation. White matter and ventricular regions were omitted entirely.

### Regional expression and similarity matrices

We created the mouse and human gene-by-region expression matrices from the mouse gene-by-voxel and human gene-by-sample expression matrices. First, we intersected the gene sets in these matrices with a list of 3331 homologous genes obtained from the NCBI HomoloGene database *(58)*, resulting in 2624 homologous genes present in both the mouse and human expression matrices. We then annotated each of the human samples with one of the 88 human atlas regions, and each of the mouse voxels with one of the 67 mouse atlas regions, discarding white matter and ventricular entries in the process. These labelled expression matrices were subsequently normalized as follows: For each matrix, we first standardized every gene across all voxels/samples using a z-scoring procedure. We then centered every voxel/sample by subtracting the average expression over all homologous genes. Finally, we generated the gene-by-region expression matrices by averaging the expression of every gene over the voxels/samples corresponding to each atlas region. Using these expression matrices, we generated the mouse-human similarity matrix by computing the Pearson correlation coefficient between all pairs of mouse and human regions.

### Multi-layer perceptron classification and latent space

To improve the resolution of mouse-human neuroanatomical matches, we performed a supervised learning approach, wherein we trained a multi-layer perceptron neural network to classify 67 mouse atlas regions from the expression values of 2624 homologous genes. We chose a model architecture in which each layer of the network was fully connected to previous and subsequent layers. To optimize the hyperparameters, we implemented an ad hoc cross-validation procedure that took into account the fact that the majority of genes in the coronal AMBA data set are sampled only once over the entire mouse brain. The procedure involved a combination of the coronal data set and the sagittal in-situ hybridization data sets. For the sagittal data set, we used the expression matrix described above. However, we used a modified version of the coronal expression matrix. This matrix was generated using the pipeline described above with the following modifications: 1. We applied a *unilateral* brain mask to the coronal images since the sagittal data is unilateral by construction, and 2. we did not aggregate the expression of multiple in-situ hybridization experiments for those genes in the coronal set pertaining to more than one experiment. We then filtered these experiment-by-voxel expression matrices according to the list of mouse-human homologous genes, as well as the human sample expression matrix. We also annotated the voxels in each of the expression matrices with one of the 67 regions in the mouse atlas. Our validation procedure then involved iterative construction of training and validation sets by sampling gene experiments from either the coronal or sagittal matrices: For every gene in the homologous set, we first determined whether that gene was associated with more than one experiment in the coronal matrix. If this was the case, we randomly sampled one of those experiments for the training set and one of the remaining experiments for the validation set. If the gene was associated with only one experiment in the coronal set, we randomly sampled either the coronal or sagittal experiment for the training set and the other for the validation set. Once the training and validation sets were generated, they were normalized using the procedure described above. We then trained the neural network using the training set and evaluated its performance on the validation set. Given that the construction of the training and validation sets involved some stochasticity, we repeated this construction, training, and validation procedure 10 times for every combination of hyperparameters.

The hyperparameters that we optimized using this method were the number of hidden layers in the network, the number of hidden units (i.e. neurons) per hidden layer, the dropout rate, and the amount weight decay. The values we optimized over were as follows. We varied the number of hidden layers between 3 and 5. We varied the number of hidden units between 100 and 1000 in increments of 100. For the dropout rate, we examined values of 0, 0.25 and 0.50. For the amount of weight decay, we examined values of 10^*−*6^ and 10^*−*5^. We found that the best-performing model had 3 hidden layers, 200 neurons per layer, a dropout rate of 0, and a weight decay value of 10^*−*6^. This model returned an average classification accuracy of 0.926 on the training sets and of 0.526 on the validation sets. Following validation, we used the optimal hyperparameters to train the network on the full bilateral coronal voxel-wise expression matrix.

These models were implemented in Python using PyTorch via the skorch library (https://skorch.readthedocs.io/en/stable/).For both validation and training, the models were trained over 200 epochs using a maximum learning rate of 10^*−*5^. We used the AdamW optimization algorithm *(59)* and OneCycleLR learning rate scheduler policy. The activation function used in the forward pass was the rectified linear unit (ReLU) and the loss function was the negative log-likelihood loss, which is the default for the NeuralNetClassifier class in skorch.

We used the trained perceptron to generate the latent gene expression space. To extract the appropriate transformation, we removed the predictive output layer and soft-max transformation from the network architecture. The resulting architecture returns the 200 hidden units in the third hidden layer as the output of the network. To create the latent space data representations, we applied this network to the mouse and human gene-by-region expression matrices, transposed so that the genes were the input variables. The resulting matrices have 200 columns corresponding to the hidden units and 67 (88) rows corresponding to the mouse (human) regions. They describe each brain region’s weight over the hidden units. To create the similarity matrix, we calculated the Pearson correlation coefficient between all pairs of mouse and human regions.

Given the stochasticity inherent in training the network (e.g. random weight initialization and stochastic optimization), we repeated the training and transformation process 500 times using the same network architecture and input data.

## Data and code availability

This manuscript, including all figures, was generated programmatically using R Markdown (https://rmarkdown.rstudio.com) and LATEX(https://www.latex-project.org). The Allen Mouse Brain Atlas and Allen Human Brain Atlas data sets are openly accessible and can be downloaded from the Allen Institute’s API (http://api.brain-map.org). All of the code and additional data needed to generate this analysis, including figures and manuscript, is accessible at https://github.com/abeaucha/MouseHumanTranscriptomicSimilarity/.

## References

1. H. J. Hedrich, H. Mossmann, W. Nicklas, in The Laboratory Mouse (Elsevier Academic Press, 1st Edition., 2004), pp. 395–408.

2. L.-M. Houdebine, in The Laboratory Mouse (Elsevier Academic Press, 1st Edition., 2004), pp. 97–107.

3. M. R. Dietrich, R. A. Ankeny, P. M. Chen, Publication Trends in Model Organism Research. Genetics. 198, 787–794 (2014).

4. B. Ellenbroek, J. Youn, Rodent models in neuroscience research: Is it a rat race? Disease Models & Mechanisms. 9, 1079–1087 (2016).

5. S. W. Oh, J. A. Harris, L. Ng, B. Winslow, N. Cain, S. Mihalas, Q. Wang, C. Lau, L. Kuan, A. M. Henry, M. T. Mortrud, B. Ouellette, T. N. Nguyen, S. A. Sorensen, C. R. Slaughterbeck, W. Wakeman, Y. Li, D. Feng, A. Ho, E. Nicholas, K. E. Hirokawa, P. Bohn, K. M. Joines, H. Peng, M. J. Hawrylycz, J. W. Phillips, J. G. Hohmann, P. Wohnoutka, C. R. Gerfen, C. Koch, A. Bernard, C. Dang, A. R. Jones, H. Zeng, A mesoscale connectome of the mouse brain. Nature. 508, 207–214 (2014).

6. R. D. Hodge, T. E. Bakken, J. A. Miller, K. A. Smith, E. R. Barkan, L. T. Graybuck, J. L. Close, B. Long, N. Johansen, O. Penn, Z. Yao, J. Eggermont, T. Höllt, B. P. Levi, S. I. Shehata, B. Aevermann, A. Beller, D. Bertagnolli, K. Brouner, T. Casper, C. Cobbs, R. Dalley, N. Dee, S.-L. Ding, R. G. Ellenbogen, O. Fong, E. Garren, J. Goldy, R. P. Gwinn, D. Hirschstein, C. D. Keene, M. Keshk, A. L. Ko, K. Lathia, A. Mahfouz, Z. Maltzer, M. McGraw, T. N. Nguyen, J. Nyhus, J. G. Ojemann, A. Oldre, S. Parry, S. Reynolds, C. Rimorin, N. V. Shapovalova, S. Somasundaram, A. Szafer, E. R. Thomsen, M. Tieu, G. Quon, R. H. Scheuermann, R. Yuste, S. M. Sunkin, B. Lelieveldt, D. Feng, L. Ng, A. Bernard, M. Hawrylycz, J. W. Phillips, B. Tasic, H. Zeng, A. R. Jones, C. Koch, E. S. Lein, Conserved cell types with divergent features in human versus mouse cortex. Nature. 573, 61–68 (2019).

7. Z. Yao, C. T. J. van Velthoven, T. N. Nguyen, J. Goldy, A. E. Sedeno-Cortes, F. Baftizadeh, D. Bertagnolli, T. Casper, M. Chiang, K. Crichton, S.-L. Ding, O. Fong, E. Garren, A. Glandon, N. W. Gouwens, J. Gray, L. T. Graybuck, M. J. Hawrylycz, D. Hirschstein, M. Kroll, K. Lathia, C. Lee, B. Levi, D. McMillen, S. Mok, T. Pham, Q. Ren, C. Rimorin, N. Shapovalova, J. Sulc, S. M. Sunkin, M. Tieu, A. Torkelson, H. Tung, K. Ward, N. Dee, K. A. Smith, B. Tasic, H. Zeng, A taxonomy of transcriptomic cell types across the isocortex and hippocampal formation. Cell. 184, 3222–3241.e26 (2021).

8. M. Hay, D. W. Thomas, J. L. Craighead, C. Economides, J. Rosenthal, Clinical development success rates for investigational drugs. Nature Biotechnology. 32, 40–51 (2014).

9. J. H. Kaas, The evolution of neocortex in primates. Progress in brain research. 195, 91–102 (2012).

10. L. Krubitzer, The Magnificent Compromise: Cortical Field Evolution in Mammals. Neuron. 56, 201–208 (2007).

11. T. M. Preuss, Do rats have prefrontal cortex? The Rose-Woolsey-Akert program reconsidered. Journal of Cognitive Neuroscience. 7, 24 (1995).

12. M. Laubach, L. M. Amarante, K. Swanson, S. R. White, What, If Anything, Is Rodent Prefrontal Cortex? eneuro. 5, ENEURO.0315–18.2018 (2018).

13. S. van Heukelum, R. B. Mars, M. Guthrie, J. K. Buitelaar, C. F. Beckmann, P. H. E. Tiesinga, B. A. Vogt, J. C. Glennon, M. N. Havenith, Where is Cingulate Cortex? A Cross-Species View. Trends in Neurosciences. 43, 285–299 (2020).

14. R. B. Mars, S. Jbabdi, M. F. S. Rushworth, A Common Space Approach to Comparative Neuroscience. Annual Review of Neuroscience. 44, 69–86 (2021).

15. R. B. Mars, S. N. Sotiropoulos, R. E. Passingham, J. Sallet, L. Verhagen, A. A. Khrapitchev, N. Sibson, S. Jbabdi, Whole brain comparative anatomy using connectivity blueprints. eLife. 7 (2018), doi:10.7554/eLife.35237.

16. R. E. Passingham, K. E. Stephan, R. Kötter, The anatomical basis of functional localization in the cortex. Nature Reviews Neuroscience. 3, 606–616 (2002).

17. R. B. Mars, R. E. Passingham, S. Jbabdi, Connectivity Fingerprints: From Areal Descriptions to Abstract Spaces. Trends in Cognitive Sciences. 22, 1026–1037 (2018).

18. R. B. Mars, L. Verhagen, T. E. Gladwin, F.-X. Neubert, J. Sallet, M. F. S. Rushworth, Comparing brains by matching connectivity profiles. Neuroscience & Biobehavioral Reviews. 60, 90–97 (2016).

19. R. B. Mars, J. Sallet, F.-X. Neubert, M. F. S. Rushworth, Connectivity profiles reveal the relationship between brain areas for social cognition in human and monkey temporoparietal cortex. Proceedings of the National Academy of Sciences. 110, 10806–10811 (2013).

20. J. Sallet, R. B. Mars, M. P. Noonan, F.-X. Neubert, S. Jbabdi, J. X. O’Reilly, N. Filippini, A. G. Thomas, M. F. Rushworth, The Organization of Dorsal Frontal Cortex in Humans and Macaques. Journal of Neuroscience. 33, 12255–12274 (2013).

21. F.-X. Neubert, R. B. Mars, A. G. Thomas, J. Sallet, M. F. S. Rushworth, Comparison of Human Ventral Frontal Cortex Areas for Cognitive Control and Language with Areas in Monkey Frontal Cortex. Neuron. 81, 700–713 (2014).

22. J. H. Balsters, V. Zerbi, J. Sallet, N. Wenderoth, R. B. Mars, Primate homologs of mouse corticostriatal circuits. eLife. 9, 24 (2020).

23. D. J. Schaeffer, Y. Hori, K. M. Gilbert, J. S. Gati, R. S. Menon, S. Everling, Divergence of rodent and primate medial frontal cortex functional connectivity. Proceedings of the National Academy of Sciences. 117, 21681–21689 (2020).

24. C. Ortiz, J. Fenandez Navarro, A. Jurek, A. Märtin, J. Lundeberg, K. Meletis, Molecular atlas of the adult mouse brain. Science Advances. 6, 14 (2020).

25. E. S. Lein, M. J. Hawrylycz, N. Ao, M. Ayres, A. Bensinger, A. Bernard, A. F. Boe, M. S. Boguski, K. S. Brockway, E. J. Byrnes, L. Chen, L. Chen, T.-M. Chen, M. Chi Chin, J. Chong, B. E. Crook, A. Czaplinska, C. N. Dang, S. Datta, N. R. Dee, A. L. Desaki, T. Desta, E. Diep, T. A. Dolbeare, M. J. Donelan, H.-W. Dong, J. G. Dougherty, B. J. Duncan, A. J. Ebbert, G. Eichele, L. K. Estin, C. Faber, B. A. Facer, R. Fields, S. R. Fischer, T. P. Fliss, C. Frensley, S. N. Gates, K. J. Glattfelder, K. R. Halverson, M. R. Hart, J. G. Hohmann, M. P. Howell, D. P. Jeung, R. A. Johnson, P. T. Karr, R. Kawal, J. M. Kidney, R. H. Knapik, C. L. Kuan, J. H. Lake, A. R. Laramee, K. D. Larsen, C. Lau, T. A. Lemon, A. J. Liang, Y. Liu, L. T. Luong, J. Michaels, J. J. Morgan, R. J. Morgan, M. T. Mortrud, N. F. Mosqueda, L. L. Ng, R. Ng, G. J. Orta, C. C. Overly, T. H. Pak, S. E. Parry, S. D. Pathak, O. C. Pearson, R. B. Puchalski, Z. L. Riley, H. R. Rockett, S. A. Rowland, J. J. Royall, M. J. Ruiz, N. R. Sarno, K. Schaffnit, N. V. Shapovalova, T. Sivisay, C. R. Slaughterbeck, S. C. Smith, K. A. Smith, B. I. Smith, A. J. Sodt, N. N. Stewart, K.-R. Stumpf, S. M. Sunkin, M. Sutram, A. Tam, C. D. Teemer, C. Thaller, C. L. Thompson, L. R. Varnam, A. Visel, R. M. Whitlock, P. E. Wohnoutka, C. K. Wolkey, V. Y. Wong, M. Wood, M. B. Yaylaoglu, R. C. Young, B. L. Youngstrom, X. Feng Yuan, B. Zhang, T. A. Zwingman, A. R. Jones, Genome-wide atlas of gene expression in the adult mouse brain. Nature. 445, 168–176 (2007).

26. M. J. Hawrylycz, E. S. Lein, A. L. Guillozet-Bongaarts, E. H. Shen, L. Ng, J. A. Miller, L. N. van de Lagemaat, K. A. Smith, A. Ebbert, Z. L. Riley, C. Abajian, C. F. Beckmann, A. Bernard, D. Bertagnolli, A. F. Boe, P. M. Cartagena, M. M. Chakravarty, M. Chapin, J. Chong, R. A. Dalley, B. D. Daly, C. Dang, S. Datta, N. Dee, T. A. Dolbeare, V. Faber, D. Feng, D. R. Fowler, J. Goldy, B. W. Gregor, Z. Haradon, D. R. Haynor, J. G. Hohmann, S. Horvath, R. E. Howard, A. Jeromin, J. M. Jochim, M. Kinnunen, C. Lau, E. T. Lazarz, C. Lee, T. A. Lemon, L. Li, Y. Li, J. A. Morris, C. C. Overly, P. D. Parker, S. E. Parry, M. Reding, J. J. Royall, J. Schulkin, P. A. Sequeira, C. R. Slaughterbeck, S. C. Smith, A. J. Sodt, S. M. Sunkin, B. E. Swanson, M. P. Vawter, D. Williams, P. Wohnoutka, H. R. Zielke, D. H. Geschwind, P. R. Hof, S. M. Smith, C. Koch, S. G. N. Grant, A. R. Jones, An anatomically comprehensive atlas of the adult human brain transcriptome. Nature. 489, 391–399 (2012).

27. J. B. Burt, M. Demirtaş, W. J. Eckner, N. M. Navejar, J. L. Ji, W. J. Martin, A. Bernacchia, A. Anticevic, J. D. Murray, Hierarchy of transcriptomic specialization across human cortex captured by structural neuroimaging topography. Nature Neuroscience. 21, 1251–1259 (2018).

28. B. D. Fulcher, J. D. Murray, V. Zerbi, X.-J. Wang, Multimodal gradients across mouse cortex. Proceedings of the National Academy of Sciences. 116, 4689–4695 (2019).

29. M. Englund, S. S. James, R. Bottom, K. J. Huffman, S. P. Wilson, L. A. Krubitzer, Comparing cortex-wide gene expression patterns between species in a common reference frame (2021), doi:10.1101/2021.07.28.454203.

30. J. H. Kaas, Reconstructing the Areal Organization of the Neocortex of the First Mammals. Brain, Behavior and Evolution. 78, 7–21 (2011).

31. G. Horev, J. Ellegood, J. P. Lerch, Y.-E. E. Son, L. Muthuswamy, H. Vogel, A. M. Krieger, A. Buja, R. M. Henkelman, M. Wigler, A. A. Mills, Dosage-dependent phenotypes in models of 16p11.2 lesions found in autism. Proceedings of the National Academy of Sciences. 108, 17076–17081 (2011).

32. A. L. Gompers, L. Su-Feher, J. Ellegood, N. A. Copping, M. A. Riyadh, T. W. Stradleigh, M. C. Pride, M. D. Schaffler, A. A. Wade, R. Catta-Preta, I. Zdilar, S. Louis, G. Kaushik, B. J. Mannion, Plajzer-Frick V. Afzal, A. Visel, L. A. Pennacchio, D. E. Dickel, J. P. Lerch, J. N. Crawley, K. S. Zarbalis, J. L. Silverman, A. S. Nord, Germline Chd8 haploinsufficiency alters brain development in mouse. Nature Neuroscience. 20, 1062–1073 (2017).

33. M. Pagani, N. Barsotti, A. Bertero, S. Trakoshis, L. Ulysse, A. Locarno, I. Miseviciute, A. De Felice, C. Canella, K. Supekar, A. Galbusera, V. Menon, R. Tonini, G. Deco, M. V. Lombardo, M. Pasqualetti, A. Gozzi, mTOR-related synaptic pathology causes autism spectrum disorder-associated functional hyperconnectivity. Nature Communications. 12, 6084 (2021).

34. L. Ventura-Antunes, B. Mota, S. Herculano-Houzel, Different scaling of white matter volume, cortical connectivity, and gyrification across rodent and primate brains. Frontiers in Neuroanatomy. 7 (2013), doi:10.3389/fnana.2013.00003.

35. P. L. Stahl, F. Salmén, S. Vickovic, A. Lundmark, J. F. Navarro, J. Magnusson, S. Giacomello, M. Asp, J. O. Westholm, M. Huss, A. Mollbrink, S. Linnarsson, S. Codeluppi, Å. Borg, F. Pontén, P. Costea, P. Sahlén, J. Mulder, O. Bergmann, J. Lundeberg, J. Frisén, Visualization and analysis of gene expression in tissue sections by spatial transcriptomics. Science. 353, 78–82 (2016).

36. S. Vickovic, G. Eraslan, F. Salmén, J. Klughammer, L. Stenbeck, D. Schapiro, T. Äijö, R. Bonneau, L. Bergenstråhle, J. F. Navarro, J. Gould, G. K. Griffin, Å. Borg, M. Ronaghi, J. Frisén, J. Lundeberg, A. Regev, P.L. Ståhl, High-definition spatial transcriptomics for in situ tissue profiling. Nature methods. 16, 987–990 (2019).

37. E. Myers, thesis, Boston University (2017).

38. G. F. Striedter, R. G. Northcutt, Brains Through Time (Oxford University Press, 2020).

39. T. A. Chaplin, H.-H. Yu, J. G. M. Soares, R. Gattass, M. G. P. Rosa, A Conserved Pattern of Differential Expansion of Cortical Areas in Simian Primates. Journal of Neuroscience. 33, 15120– 15125 (2013).

40. R. B. Mars, R. E. Passingham, F.-X. Neubert, L. Verhagen, J. Sallet, in Evolution of Nervous Systems (Academic Press, 2016), vol. 4, xp. 185–205.

41. N. Eichert, E. C. Robinson, K. L. Bryant, S. Jbabdi, M. Jenkinson, L. Li, K. Krug, K. E. Watkins, R. B. Mars, Cross-species cortical alignment identifies different types of anatomical reorganization in the primate temporal lobe. eLife. 9, e53232 (2020).

42. P. H. Rudebeck, A. Izquierdo, Foraging with the frontal cortex: A cross-species evaluation of reward-guided behavior. Neuropsychopharmacology. 47, 134–146 (2022).

43. J. H. Kaas, Neocortex in early mammals and its subsequent variations. Annals of the New York Academy of Sciences. 1225, 10.1111/j.1749–6632.2011.05981.x (2011).

44. H. C. Barron, R. B. Mars, D. Dupret, J. P. Lerch, C. Sampaio-Baptista, Cross-species neuroscience: Closing the explanatory gap. Philosophical Transactions of the Royal Society B: Biological Sciences. 376, 20190633 (2021).

45. X. Liu, S. B. Eickhoff, S. Caspers, J. Wu, S. Genon, F. Hoffstaedter, R. B. Mars, I. E. Sommer, C. R. Eickhoff, J. Chen, R. Jardri, K. Reetz, I. Dogan, A. Aleman, L. Kogler, O. Gruber, J. Caspers, C. Mathys, K. R. Patil, Functional parcellation of human and macaque striatum reveals human-specific connectivity in the dorsal caudate. NeuroImage. 235, 118006 (2021).

46. F. Mandino, R. M. Vrooman, H. E. Foo, L. Y. Yeow, T. A. W. Bolton, P. Salvan, C. L. Teoh, C. Y. Lee, A. Beauchamp, S. Luo, R. Bi, J. Zhang, G. H. T. Lim, N. Low, J. Sallet, J. Gigg, J. P. Lerch, R. B. Mars, M. Olivo, Y. Fu, J. Grandjean, A triple-network organization for the mouse brain. Molecular Psychiatry, 1–8 (2021).

47. E. Simonoff, A. Pickles, T. Charman, S. Chandler, T. Loucas, G. Baird, Psychiatric Disorders in Children With Autism Spectrum Disorders: Prevalence, Comorbidity, and Associated Factors in a Population-Derived Sample. Journal of the American Academy of Child & Adolescent Psychiatry. 47, 921–929 (2008).

48. R. Grzadzinski, M. Huerta, C. Lord, DSM-5 and autism spectrum disorders (ASDs): An opportunity for identifying ASD subtypes. Molecular Autism. 4, 12 (2013).

49. J. Ellegood, E. Anagnostou, B. A. Babineau, J. N. Crawley, L. Lin, M. Genestine, E. DiCicco-Bloom, J. K. Y. Lai, J. A. Foster, O. Peñagarikano, D. H. Geschwind, L. K. Pacey, D. R. Hampson, C.L. Laliberté, A. A. Mills, E. Tam, L. R. Osborne, M. Kouser, F. Espinosa-Becerra, Z. Xuan, C. M. Powell, A. Raznahan, D. M. Robins, N. Nakai, J. Nakatani, T. Takumi, M. C. van Eede, T. M. Kerr, C. Muller, R. D. Blakely, J. Veenstra-VanderWeele, R. M. Henkelman, J. P. Lerch, Clustering autism: Using neuroanatomical differences in 26 mouse models to gain insight into the heterogeneity. Molecular Psychiatry. 20, 118–125 (2015).

50. Q. Wang, S.-L. Ding, Y. Li, J. Royall, D. Feng, P. Lesnar, N. Graddis, M. Naeemi, B. Facer, A. Ho, T. Dolbeare, B. Blanchard, N. Dee, W. Wakeman, K. E. Hirokawa, A. Szafer, S. M. Sunkin, S. W. Oh, A. Bernard, J. W. Phillips, M. Hawrylycz, C. Koch, H. Zeng, J. A. Harris, L. Ng, The Allen Mouse Brain Common Coordinate Framework: A 3D Reference Atlas. Cell. 181, 936–953.e20 (2020).

51. R. D. Markello, A. Arnatkeviciūtė, J.-B. Poline, B. D. Fulcher, A. Fornito, Standardizing workflows in imaging transcriptomics with the abagen toolbox. Biorxiv, 22 (2021).

52. A. Arnatkeviciūtė, B. D. Fulcher, A. Fornito, A practical guide to linking brain-wide gene expression and neuroimaging data. NeuroImage. 189, 353–367 (2019).

53. A. E. Dorr, J. P. Lerch, S. Spring, N. Kabani, R. M. Henkelman, High resolution three-dimensional brain atlas using an average magnetic resonance image of 40 adult C57Bl/6J mice. NeuroImage. 42, 60–69 (2008).

54. K. Richards, C. Watson, R. F. Buckley, N. D. Kurniawan, Z. Yang, M. D. Keller, R. Beare, P. F. Bartlett, G. F. Egan, G. J. Galloway, G. Paxinos, S. Petrou, D. C. Reutens, Segmentation of the mouse hippocampal formation in magnetic resonance images. NeuroImage. 58, 732–740 (2011).

55. J. F. P. Ullmann, C. Watson, A. L. Janke, N. D. Kurniawan, D. C. Reutens, A segmentation protocol and MRI atlas of the C57BL/6J mouse neocortex. NeuroImage. 78, 196–203 (2013).

56. P. E. Steadman, J. Ellegood, K. U. Szulc, D. H. Turnbull, A. L. Joyner, R. M. Henkelman, J. P. Lerch, Genetic Effects on Cerebellar Structure Across Mouse Models of Autism Using a Magnetic Resonance Imaging Atlas: MRI of genetic mouse model’s cerebellum. Autism Research. 7, 124–137 (2014).

57. L. R. Qiu, D. J. Fernandes, K. U. Szulc-Lerch, J. Dazai, B. J. Nieman, D. H. Turnbull, J. A. Foster, M. R. Palmert, J. P. Lerch, Mouse MRI shows brain areas relatively larger in males emerge before those larger in females. Nature Communications. 9, 2615 (2018).

58. NCBI, Database resources of the National Center for Biotechnology Information. Nucleic Acids Research. 46, D8–D13 (2018).

59. I. Loshchilov, F. Hutter, in Proceedings of the Seventh International Conference on Learning Representations (New Orleans, 2019; http://arxiv.org/abs/1711.05101).

